# Primitive Hepatoblasts Driving Early Liver Development

**DOI:** 10.1101/2025.06.08.658502

**Authors:** Kentaro Iwasawa, Hiroyuki Koike, Hasan Al Reza, Yuka Milton, Keishi Kishimoto, Konrad Thorner, Marissa Granitto, Norikazu Saiki, Connie Santangelo, Kathryn Glaser, Masaki Kimura, Alexander Bondoc, Hee-Wong Lim, Mitsuru Morimoto, Makiko Iwafuchi, James M. Wells, Aaron M. Zorn, Takanori Takebe

## Abstract

The embryonic development of the liver is initiated by the emergence of hepatoblasts, originating from the ventral foregut endoderm adjacent to the heart. Here, we identify and characterize a previously unrecognized population of early hepatoblasts at the ventroposterior part of the emerging liver bud, traced from *Cdx2*-positive endoderm progenitors, which we term primitive hepatoblasts. Mouse and human single-cell transcriptomics reveals the expression of both canonical hepatoblast markers *TBX3*, *FGB*, and *KRT8/18* and primitive-specific mesenchymal markers *ID3*, *VIM*, and *GATA4*. Lineage tracing revealed the notable contribution up to 12.6% of LIV2+ hepatoblasts at E11.5 but diminishes in late fetal and postnatal development. Epigenetic and functional perturbation studies further uncover that primitive hepatoblast emergence is primed by WNT-suppression on RA-permissive CDX2+FOXA2+ progenitors. Furthermore, human pluripotent stem cell-derived primitive hepatoblast-like cells secrete pleiotrophin and midkine to amplify hepatoblast populations and develop epithelial-mesenchymal hybrid tissues *in vivo*. Our results provide a new framework for understanding lineage heterogeneity during early hepatogenesis and offer revised insights into strategies to model normal and abnormal liver development.

## Introduction

The embryonic development of the liver bud is initiated by the emergence of hepatoblasts, bipotent progenitors that drive exponential tissue growth during early development. While directed differentiation efforts in guiding human pluripotent stem cells (hPSCs) into hepatic lineages have advanced extensively, current protocols fail to capture the transient amplification program in part due to a lack of full recapitulation of intercellular communication during this critical stage.

Developmental biology studies have described distinct subpopulations of hepatoblasts with transcriptional and phenotypic differences and suggest that early hepatoblasts are not a homogeneous population^1–5^. Supporting the concept of early lineage diversification, fate-mapping studies in mice have shown that progenitor cells located around the anterior intestinal portal (AIP) lip at embryonic day (E) 8.0–8.5 give rise to both the liver bud and ventral midgut later in development^6^. This study identified two spatially distinct endodermal progenitor populations, indicating a heterogeneous origin of hepatoblasts as early as this stage. Single-cell transcriptomics has revealed multiple layers of hepatoblast heterogeneity across species. Lgr5 hepatoblasts exhibit bipotent differentiation potential with unique molecular signatures^1^ and pancreato-biliary progenitors have been shown to contribute to the expanding hepatoblast pool during liver bud outgrowth^2^. Moreover, subpopulations such as hepatomesenchyme^3^ and ID3 hepatoblasts^4^ co-express mesenchymal markers^3^, while retaining comparable hepatocyte differentiation capacity^4^. Similarly, lineage-tracing studies in zebrafish have identified a mixture of unipotent and bipotent hepatoblasts^7^. However, the cellular and functional heterogeneity among hepatoblasts remains incompletely understood. A deeper understanding of the lineage trajectories and roles of these heterogeneous hepatoblasts is essential for navigating in vitro strategies to dictate liver bud development.

Prospective hepatic domain emerges at the foregut–midgut boundary. SOX2 marks anterior foregut identity, while CDX2 is a hallmark of mid/hindgut fate ^8,9^. FOXA pioneer transcription factors (TFs) including FOXA2 broadly establish chromatin accessibility at hepatic enhancers^10^ in the developing liver^11^. These TF expressions exhibit redundant expression patterns in the early endoderm, albeit preserving hepatic developmental potential from either gene knockout^12^. To uncover how these transcriptionally distinct progenitor populations contribute to the emergence and heterogeneity of hepatoblasts, we herein employed lineage tracing by *Foxa2, Cdx2* and *Sox2*, high-resolution imaging, single cell profiling and stem cell differentiation approaches to investigate the hepatoblast heterogeneity during early liver development. We identified a previously unrecognized class of hepatoblasts, which we term "primitive hepatoblasts." These cells are derived from Cdx2-positive putative endodermal progenitors and exhibit distinct molecular and functional characteristics compared to other hepatoblast subpopulations, suggesting a unique role in liver development.

## Results

### Spatial characterization of Cdx2 expression during early liver development

To test the hypothesis that CDX2-positive endodermal cells might contribute to the embryonic liver, we conducted CDX2 wholemount immunofluorescence staining of E8.5 mouse embryos (6-somite stage). Expression was detected not only in the midgut and visceral endoderm, as expected, but also notably in the lateral domain of the anterior intestinal portal (AIP) lip endoderm (**Fig. 1a**). Importantly, these CDX2⁺ cells co-expressed PROX1, a marker of hepato-biliary-pancreatic progenitors at the location previously identified as lateral endoderm progenitors (LEPs)^6^. In contrast, SOX2 was localized cranially to the heart and was absent from the AIP lip (**Extended Data Fig.1a**). By E8.75, the liver bud had begun to form at the interface between SOX2⁺ and CDX2⁺ domains of the gut tube. CDX2⁺ cells were also present in the caudal region of the developing liver bud at this stage (**Extended Data Fig. 1b**). By E9.5, CDX2⁺ cells were restricted to both the ventral and dorsal pancreatic buds but were no longer detectable in any of hepatobiliary region at E10.5 (**Extended Data Fig. 1c**).

**Figure 1.**
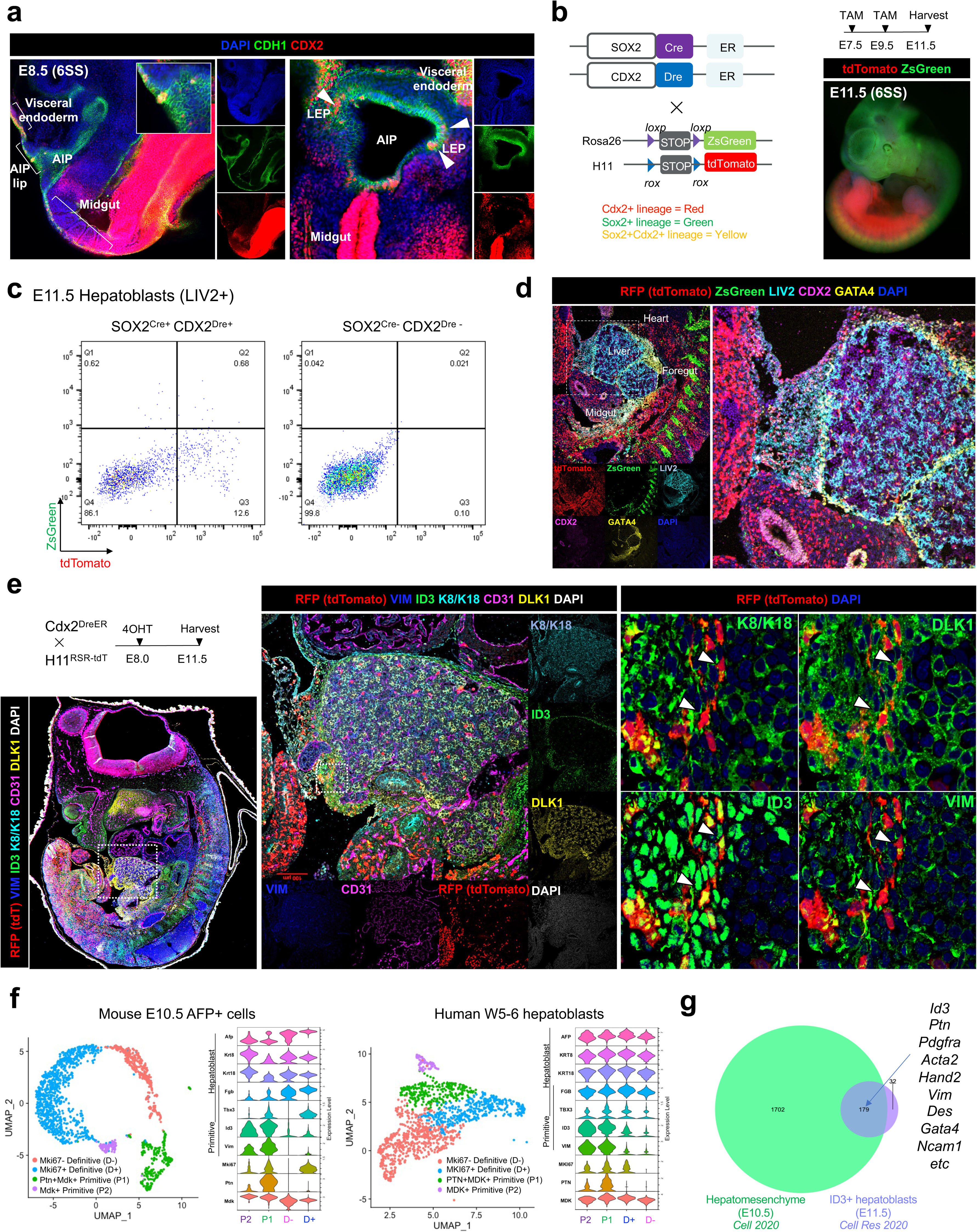
Primitive hepatoblasts in the ventroposterior edge of the developing liver co-expressing mesenchyme markers. **a.** Wholemount immunostaining of E8.5 embryo showed CDX2 expression in midgut and lateral endoderm progenitor (LEP) of the anterior intestinal portal lip. **b.** Schematic of the dual-lineage tracing strategy using Sox2^CreER^ and Cdx2^DreER^ mice crossed with dual-reporter alleles Rosa26^LSL-ZsGreen^ and H11^RSR-tdTomato^. **c.** Flow cytometry analysis of Sox2^CreER^ and Cdx2^DreER^ dual lineage-traced embryo at E11.5, gated on LIV2+ hepatoblasts. **d.** Immunostaining of E11.5 embryos revealed tdTomato+ cells within LIV2+ hepatoblasts, indicating early contribution from the Cdx2 lineage. **e.** Low-magnification (left) and high-magnification (right) immunofluorescence images at E11.5 show tdTomato-positive cells co-expressing hepatoblast markers TBX3, DLK1, KRT8/18, and ID3, along with the mesenchymal marker VIM. **f.** scRNA-seq analysis of mouse E10.5 liver (left) and human embryonic liver at weeks 5–6 (right) identified four distinct hepatoblast clusters per species. Two clusters (P1 and P2) exhibited primitive hepatoblast signatures with high *Id3/ID3* and *Vim/VIM* expression; P1 showed enriched *Ptn/PTN* expression compared to P2. Definitive endoderm clusters were separated by differential *Mki67/MKI67* expression into proliferative (D⁺) and non-proliferative (D⁻) subtypes. **g.** Venn diagram showing overlap of differentially expressed genes (DEGs) between previously reported hepatomesenchyme (Cell, 2020) and ID3⁺ hepatoblasts (Cell Research, 2020).

### Lineage tracing of hepatic progenitors from Cdx2 lineage

To trace the lineage of the early CDX2-positive cells, we employed tamoxifen inducible Dre-rox systems in mice by introducing a knock-in Cdx2^DreER^ mouse with P2A-DreERT2 cassette before the stop codon of the Cdx2 gene (**Extended Data Fig.2a**). Subsequently, we traced the lineage of Sox2 (foregut), Cdx2 (lateral AIP and midgut), and FoxA2 (gut tube) endoderm cells by crossing Sox2^CreER^, Cdx2^DreER^, or Foxa2^CreER^ mice with Rosa26^LSLtdT^ or H11^RSRtdT^ mice, followed by exposure to tamoxifen during gastrulation. Confocal imaging confirmed tamoxifen-dependent Cre- or Dre-mediated tdTomato expression in SOX2, CDX2, or FOXA2 positive cells (**Extended Data Fig.2b**). Analysis of whole-mount stained E10.5 embryos revealed a significant contribution of tdTomato-positive cells within the liver in embryos resulting from crosses between Cdx2^DreER^ and FoxA2^CreER^ lines (**Extended Data Fig.2b**).

To further investigate the progenitor populations involved in hepatic specification, we generated embryos carrying heterozygous Sox2^CreER^/Cdx2^DreER^ alleles and homozygous Rosa26^LSLZsG^/H11^RSRtdT^ alleles (**Fig. 1b**). Tamoxifen administration at E7.5 and E9.5 resulted in tdTomato labeling in 12.6% of LIV2 positive hepatoblasts at E11.5 (**Fig. 1c and 1d)** indicating a significant contribution from the Cdx2 lineage.

Subsequent Dre-rox–mediated induction revealed peak recombination efficiency at E8.0. The resulting tdTomato-positive cells expressed hepatoblast markers such as TBX3, FGB, DLK1, KRT8/18, E-cadherin and ID3, as well as the mesenchymal markers VIM and GATA4 (**Fig 1d, 1e and Extended Data Fig. 2c**). These cells intermingling with the septum mesenchyme at E9.5 were localized at the ventroposterior edge of the liver bud surface at E11.5 and co-expressed both hepatoblast and mesenchymal markers with - 30% of the population labeled with tdTomato.

### Identification of Primitive and Definitive hepatoblast populations in human and mouse embryos

To investigate the molecular heterogeneity of early hepatoblasts, we analyzed publicly available single-cell RNA sequencing datasets from human embryos at weeks 5–6 and mouse embryos at E10. In both species, four distinct hepatoblast clusters were identified (**Fig. 1f).** Of these clusters in each dataset exhibited elevated expression of mesenchymal markers including *PTN, MDK, VIM,* and *ID3* alongside canonical hepatoblast markers such as *AFP, KRT8, KRT18, FGB,* and *TBX3.* These mesenchymal-enriched clusters were designated as *primitive hepatoblasts*. In contrast, the remaining two clusters showed lower expression of these mesenchymal genes and were classified as *definitive hepatoblasts*, hereafter. One of the definitive clusters also exhibited a high *MKI67* expression, indicating a proliferative subset (**Fig. 1f**).

To further contextualize these populations, we compared the differentially expressed genes (DEGs) from two previously described hepatoblast subtypes: hepatomesenchyme^3^ and ID3⁺ hepatoblasts^4^. These groups shared 179 overlapping DEGs, including *PTN, PDGFRA, HAND2, VIM, DES, NCAM1,* and *ID3*, reinforcing the presence of a conserved mesenchymal gene signature (**Fig. 1g**).

Species-specific differences were observed in the expression of PROX1, a key hepatic transcription factor. In mouse embryos, PROX1 expression was markedly reduced in primitive hepatoblast clusters relative to definitive ones. However, in human embryos, primitive and definitive clusters expressed PROX1 at comparable levels, suggesting interspecies differences in hepatoblast maturation or identity (**Fig. 1f**). Together, these results support the existence of primitive and definitive hepatoblast populations during early liver development, distinguished by their mesenchymal molecular profiles.

### Transient contribution of Cdx2**⁺** lineage to hepatoblast and hepatocyte populations during development

To define the temporal window during which Cdx2-derived cells contribute to the fetal liver bud, we performed Dre-rox–mediated lineage tracing by induction at sequential embryonic stages (E7.5–E10.5) and analyzed embryos at E11.5. This approach revealed that tamoxifen induction at E8.0 yielded the highest labeling efficiency of tdTomato+ primitive hepatoblasts (**Fig. 2a)**.

**Figure 2.**
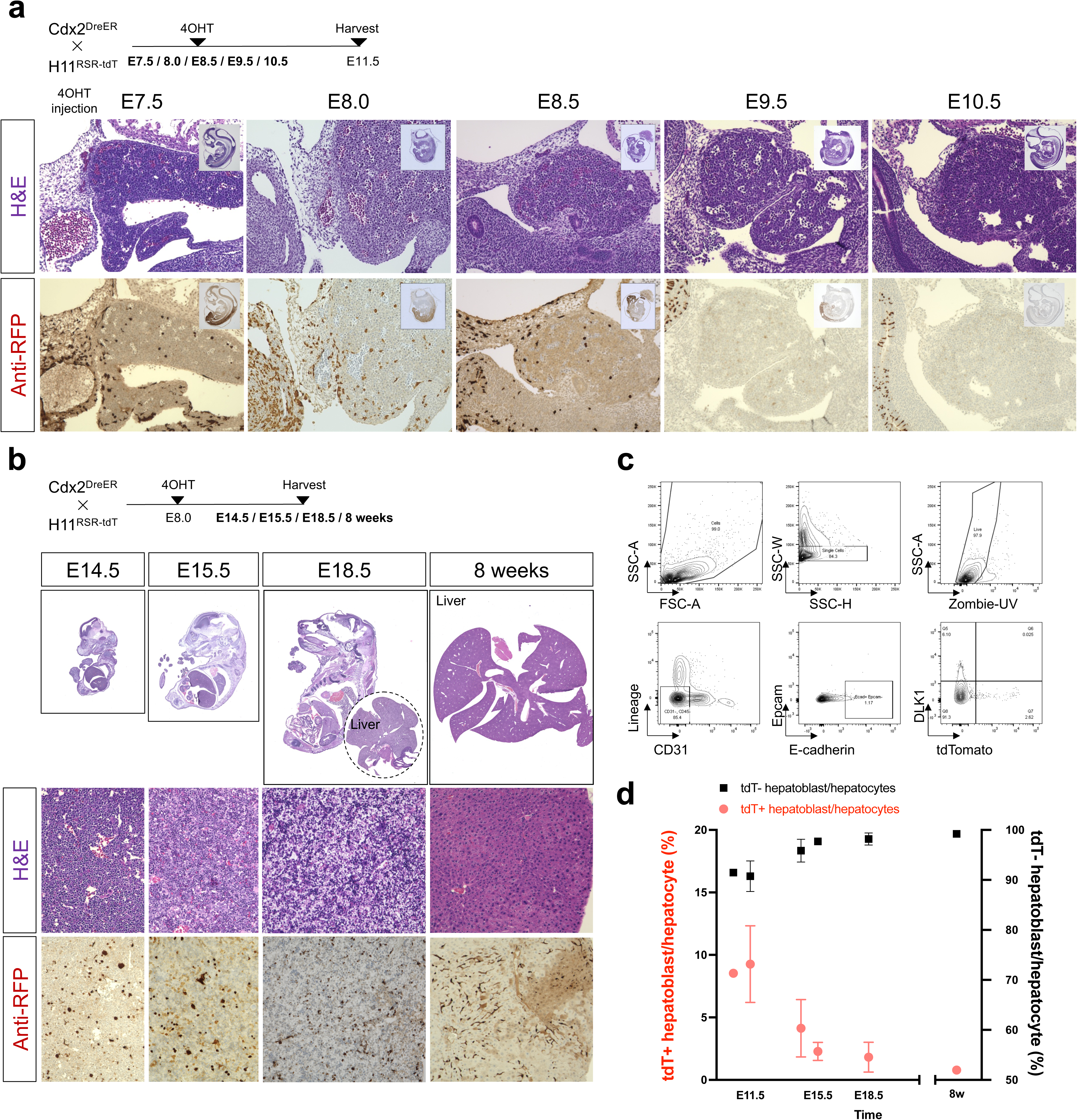
Temporal fate of Cdx2-derived primitive hepatoblasts during liver development. **a.** Dre-rox–mediated lineage tracing with 4OHT induction from E7.5 to E10.5, followed by analysis at E11.5, revealed peak recombination of Cdx2-derived hepatoblasts upon induction at E8.0. **b.** Time-course analysis of tdTomato⁺ hepatoblasts by histology from E14.5 to 8 weeks postnatally after E8.0 induction. **c.** Flow cytometry gating strategy at E18.5 revealed that ∼2.5% of E-cadherin⁺ EpCAM⁻ hepatocytes were tdTomato⁺ and lacked DLK1 expression, indicating maturation. **d.** Quantification of tdTomato⁺ hepatoblasts/hepatocytes: cells were gated as E-cadherin⁺ DLK1⁺ until E15.5, E-cadherin⁺ EpCAM⁻ at E18.5, and ASGR⁺ hepatocytes at 8 weeks. The Cdx2-derived hepatoblast population decreased to ∼3.0% by E14.5, ∼2.5% by E18.5, and ∼1% of adult hepatocytes retained tdTomato labeling at 8 weeks.

Intriguingly, this Cdx2-derived hepatoblast population declined over time. At E14.5, ∼3.0% of DLK1+E-cadherin+ hepatoblasts were tdTomato+ (**Fig. 2b**). In contrast, the proportion of tdTomato-positive endothelial and hematopoietic cells increased to 12.5% and 26.8%, respectively (**Extended Fig 3a**). By E18.5, tdTomato+ hepatocytes (E-cadherin+EpCAM-) were evenly distributed across all liver lobes, comprising approximately 2.5% of the total hepatocyte population, by which time DLK1 expression had largely diminished (**Fig. 2c, Extended Fig 3b**). In 8-week-old postnatal mice, hepatocytes were isolated using liver perfusion with digestive enzymes followed by gradient separation, revealing that ∼1% of ASGR+ hepatocytes remained tdTomato^+^ (**Extended Fig 3c**). Collectively, these lineage-tracing results indicate a transient developmental contribution from a subset of Cdx2-derived hepatoblasts, with their descendants most enriched at E11.5 (**Fig. 2d**).

### *In vitro* human primitive hepatoblast specification model

To investigate the contribution of primitive hepatoblast in humans, we modified a human PSC-derived w foregut–midgut boundary organoid model. In this system, human PSCs were differentiated into posterior gut-like (CDX2^high^; PG) and anterior gut-like (SOX2^high^; AG) populations through BMP inhibition for AG induction and WNT/FGF4 activation for both AG and PG (**Fig. 3a**). Fusion of these two populations leads to the emergence of hepato-biliary-pancreatic (HBP) progenitors at the boundary^8^. Wholemount immunofluorescence of day 11 (D11) boundary organoids generated with *PROX1::mScarlet* ESH1 cells confirmed the presence of HBP progenitors (**Fig. 3b**, **Extended Data Fig 4a and 4b**). Co-expression of PDX1 identified pancreatic progenitors, while the absence of PDX1 distinguished hepatoblast progenitors (**Fig. 3b**). Continued culture of the boundary organoids revealed an upregulation of both definitive and primitive hepatoblast progenitor markers in the CDX2^high^ PG population compared to the SOX2^high^ AG, as demonstrated by gene set enrichment analysis (**Fig. 3c and 3d**). To trace the emergence of hepatic progenitors, we next utilized the *PROX1::mScarlet* hPSC reporter line in combination with a chemical lineage-tracing strategy based on fluorescent nanoparticle-conjugated peptides^13^. This approach enables efficient labeling of individual spheroids with a success rate of 94.3%^13^. Quantitative analysis revealed significantly higher *PROX1::mScarlet* expression in the PG region compared to the AG, consistent with patterns observed using Dre-rox-based genetic tracing in mice (**Fig. 3e**).

**Figure 3.**
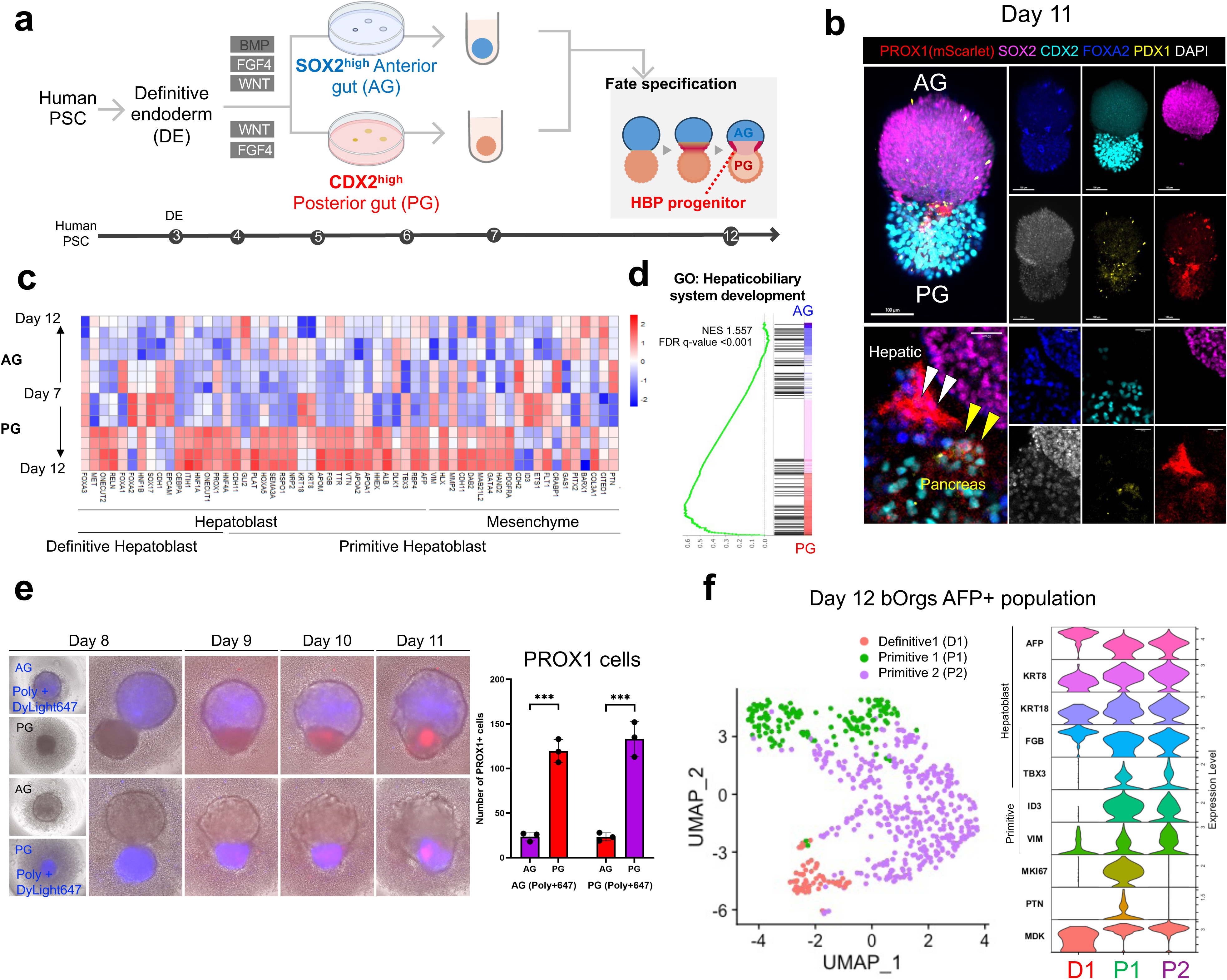
Posterior gut identity promotes primitive hepatoblast specification in boundary organoids model. **a.** Schematic of the differentiation strategy to generate anterior gut (AG) and posterior gut (PG) lineages from hPSCs, followed by fusion to model the foregut–midgut boundary. **b.** Day 11 boundary organoids derived from *PROX1::mScarlet* hPSCs exhibited both PROX1+PDX-hepatic progenitors and PROX1+PDX1+ pancreatic progenitors. Immunofluorescence markers: PROX1::mScarlet (endogenous reporter), SOX2 (magenta), CDX2 (cyan), DAPI (white), PDX1 (yellow), FOXA2 (blue). **c.** Bulk RNA sequencing revealed increased expression of primitive and definitive hepatoblast markers in PG compared to AG. **d.** Gene set enrichment analysis showed significant enrichment of hepaticobiliary system development pathways in PG versus AG. **e.** Lineage tracing using nanoparticle-mediated DyLight647 labeling in ESH1-PROX1::mScarlet-derived boundary organoids revealed the emergence of PROX1⁺ cells from the PG region. **f.** scRNA-seq of day 12 boundary organoids revealed primitive-like clusters characterized by *ID3* and *VIM* expression, alongside a smaller definitive-like cluster.

To investigate the molecular heterogeneity of early hepatoblast-like populations, we performed scRNA-seq on D12 boundary organoids (**Extended Data Fig. 5a)**. This analysis identified two major clusters of primitive hepatoblast-like cells marked by high expression of *ID3* and *VIM*, alongside canonical hepatoblast markers including *AFP, KRT8, KRT18, FGB*, and *TBX3*. In addition, we detected a smaller population resembling definitive hepatoblasts, characterized by the absence of *ID3* and *PTN* expression (**Fig. 3f**).

### CDX2+ progenitors are competent for the hepatic transcriptional program

To determine if these CDX2+ progenitors can execute the hepatic transcriptional program, we conducted CUT&RUN sequencing on both the AG and PG at D7, focusing on two crucial histone modifications: the activating marker H3K4me3 and the repressive marker H3K27me3. Our analysis identified a cumulative total of 78,148 peaks for H3K4me3 and 69,945 peaks for H3K27me3. To further link these H3K4me3 and H3K27me3 peaks with potential target genes, we leveraged the Genomic Regions Enrichment of Annotations Tool (GREAT). This bioinformatics analysis unveiled 4,033 genes associated with 6,298 AG-specific peaks and 4,789 genes linked to 6,976 PG-specific peaks for H3K27me3. Furthermore, 4,033 genes were associated with 11,681 AG-specific H3K4me3 peaks and 4,789 genes were linked to 10,639 PG-specific H3K4me3 peaks. In D7 AG cells, AG-specific gene loci were primarily marked by H3K4me3, while PG-specific loci showed both H3K4me3 and H3K27me3, indicating a poised or repressed epigenetic state. Conversely, in PG cells, PG-specific genes were enriched for H3K4me3, and AG-specific genes carried both marks. (**Fig. 4a**). To characterize the epigenetic signature at the key marker genes in the primitive hepatoblast population, we analyzed the enrichment of H3K4me3 and H3K27me3 marks in D7 AG and PG cells. Several key primitive hepatoblast-defining genes, including AFP, TBX3, and GATA4, represented poised or repressed epigenetic features in AG, whereas KRT18, TBX3, and ID3 showed an active epigenetic state in PG (**Fig. 4b**). These differential epigenetic signatures suggest transcriptional priming of the primitive hepatic program specifically in CDX2^high^ PG cells.

**Figure 4.**
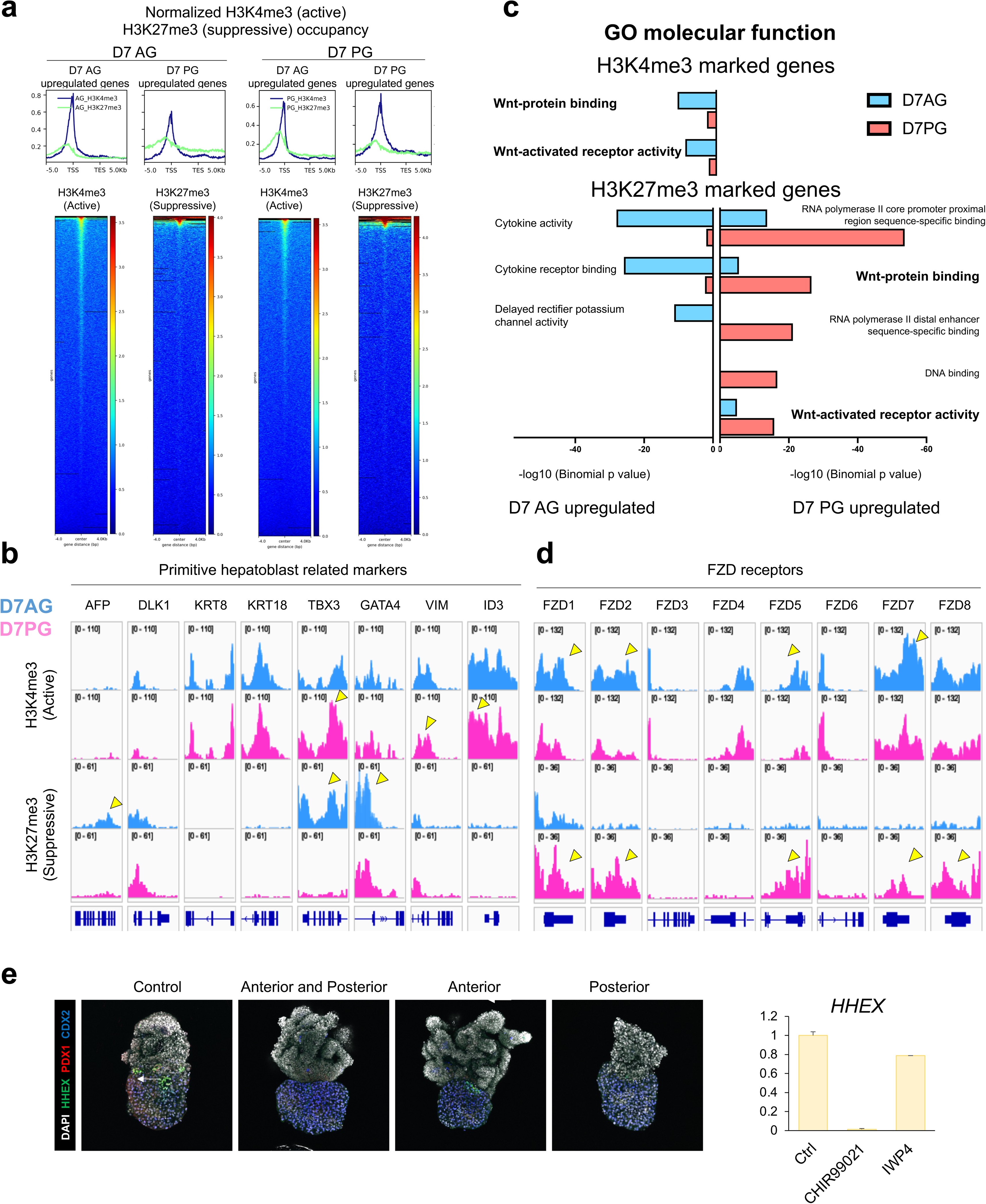
Epigenetic priming of primitive hepatoblast identity in posterior gut. **a.** Normalized enrichment scores of H3K4me3 (active mark) and H3K27me3 (suppressive mark) at D7AG- and D7PG-upregulated gene loci, confirming region-specific epigenetic priming. **b.** Primitive hepatoblast-associated genes such as TBX3 and GATA4 were marked by H3K27me3 (suppressive) enrichment in AG, while PG exhibited H3K4me3 enrichment (active) at genes including ID3, VIM, and TBX3. **c.** Gene ontology (GO) analysis revealed suppression of Wnt signaling in PG, supported by increased H3K4me3 (active) enrichment at Wnt pathway activators in AG and H3K27me3 (suppressive) enrichment at Wnt pathway activators in PG. **d.** Members of the FZD family of Wnt receptors (FZD1, FZD2, FZD5, and FZD7) showed active chromatin (H3K4me3) in AG, but were repressed (H3K27me3) in PG, consistent with epigenetic suppression of Wnt signaling in the posterior gut. **e.** Pharmacological activation of Wnt signaling using CHIR99021 impaired HHEX induction specifically when activating Wnt in PG, suggesting that Wnt inhibition is required for HBP specification.

Gene Ontology (GO) enrichment analysis revealed an upregulation of the " response to retinoic acid" pathway for H3K27me3-marked genes in AG, suggesting a reduced response to the retinoic acid (RA) in the AG as opposed to PG (**Extended Data Fig 6a**). This aligns with our prior work where AG-mesenchyme derived RA ligands activate the PG-endoderm for hepato-biliary-pancreatic specification^8^, owing to RA sensitivity in PG-endoderm. Additionally, GO terms such as "Wnt-protein binding" and "Wnt-activated receptor activity" were enriched among the H3K4me3-marked genes in AG, but among H3K27me3-marked genes in PG, underscoring a suppressed response to the Wnt pathway in PG compared to AG (**Fig. 4c**). This observation was further supported by analysis of WNT receptor gene loci, including Frizzled class receptors 1 (FZD1), FZD2, FZD5, FZD7, and FZD8, which showed higher H3K4me3 enrichment in AG and greater H3K27me3 enrichment in PG (**Fig. 4b**).

To validate the findings from trajectory analysis, we performed Wnt signaling modulation in our organoid models. Consistent with our hypothesis, activation of Wnt signaling using CHIR99021 in the PG suppressed the emergence of HHEX in the boundary organoids (**Fig. 4d**), providing further evidence for the regulatory role of Wnt signaling in HBP development^14^. These findings collectively suggest that CDX2^high^ PG endoderm are particularly sensitive to Wnt inhibition, while persistent Wnt activation impairs HBP progenitor specification at the foregut–midgut boundary.

### Enrichment of definitive hepatoblast progenitors shape hepatoblast subtype development

To investigate the impact of primitive lineages on definitive-like hepatoblast populations, we analyzed progenitor composition across timepoints. Flow cytometry (FACS) analysis shows 98−99% of AG/PG cells expresses endoderm marker FOXA2+ from D3 thru D7. At D5, AG cells were enriched for ∼45% FOXA2^+^SOX2⁻CDX2⁻ presumptive definitive hepatoblast progenitors, while D7 AG cells were exclusively FOXA2^+^SOX2^+^CDX2⁻ foregut progenitors. By using PG cells containing ∼80% FOXA2^+^SOX2⁻CDX2^+^ primitive hepatoblast progenitors, boundary organoids formed with D5 PG spheroids showed notable increase in *PROX1::mScarlet* signals by day 14. Further FACS based quantification demonstrated that ∼35% of FOXA2^+^ cells express *mScarlet*, compared to ∼20% when using D7 AG for boundary formation. These findings indicate that the coexistence of FOXA2^+^SOX2⁻CDX2⁻ and FOXA2^-^SOX2⁻CDX2+ hepatoblast progenitors orchestrates hepatic lineage development (**Fig. 5a**).

**Figure 5.**
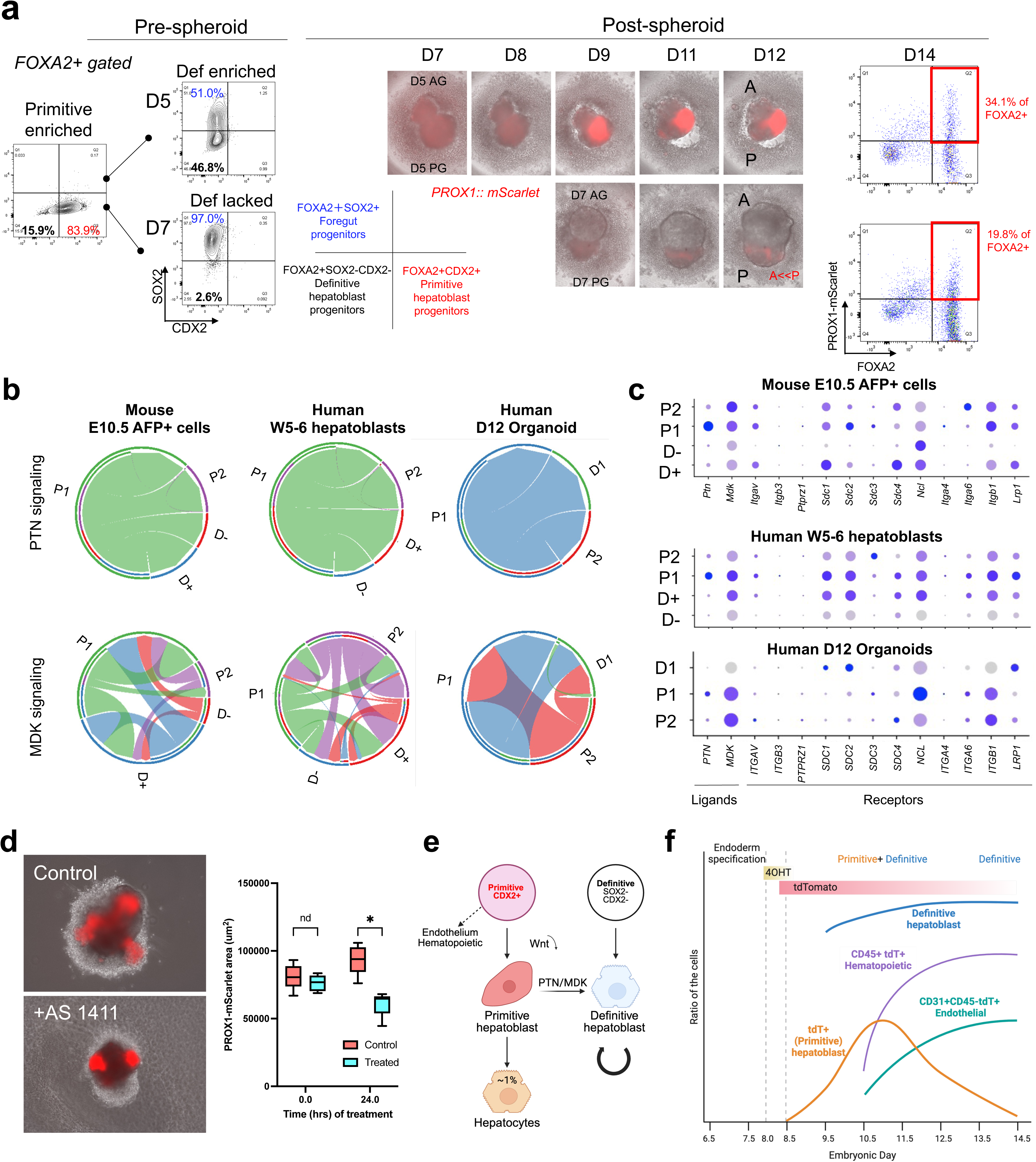
Primitive lineage-derived Pleiotrophin and Midkine signaling mediates hepatoblasts expansion. **a.** Quantification of progenitor populations pre-spheroid formation, including FOXA2+SOX2+ foregut progenitors, FOXA2+SOX2–CDX2– definitive hepatoblast progenitors, and FOXA2+CDX2+ primitive hepatoblast progenitors, at D5 and D7 in AG and PG cultures. Fusing day 5 (D5) AG and PG spheroids enhanced hepatic induction, with 34.1% of FOXA2+ cells expressing *PROX1::mScarlet* by D14, compared to 19.8% from D7 AG–PG fusions. **b.** CellChat analysis of scRNA-seq from mouse embryonic liver, human fetal liver and boundary organoids identified *Ptn/PTN* and *Mdk/MDK* signaling pathways enriched in primitive and definitive cross talk. **c.** PTN was predominantly expressed in the P1 primitive cluster, while both P1 and P2 clusters expressed MDK. Receptors such as *Ncl/NCL* were enriched in the definitive hepatoblast clusters suggesting paracrine signaling between hepatoblast subtypes. **d.** Inhibition of PTN/MDK signaling using the NCL-targeting aptamer AS1411 significantly reduced the area of PROX1+ area, demonstrating that this signaling axis is essential for progenitor expansion. **e.** Schematic illustrating the proposed lineage relationships between primitive hepatoblast progenitors (CDX2+) and definitive hepatoblast progenitors (SOX2–CDX2–) during early hepatic specification. **f.** Diagram depicting temporal changes in the abundance of primitive and definitive hepatoblast populations, alongside CD45+ hematopoietic cells and CD31+CD45– endothelial cells, highlighting dynamic shifts in cell composition during organoid development.

### Primitive hepatoblasts drive hepatoblast expansion via PTN/MDK signaling

To determine the role of primitive lineage in promoting hepatoblast expansion, we employed the CellChat computational framework to analyze scRNA-seq datasets from both mouse embryonic, human fetal hepatoblasts and human foregut-midgut boundary organoids. This analysis revealed that PTN/MDK signaling are key pathways mediating cell–cell communication between these populations (**Fig. 5b**). Primitive hepatoblast clusters expressed similar levels of *MDK*. However, the P1 cluster showed significantly higher *PTN* levels, while the P2 cluster displayed lower *PTN* expression in all three datasets. Among the two definitive hepatoblast clusters, the D+ cluster, characterized by higher *Mki67* expression, was identified as the main recipient of these signals. This cluster also exhibited high expression of PTN/MDK signaling receptors such as *NCL* (nucleolin) (**Fig. 5c**). These results suggest a PTN/MDK-mediated signaling axis facilitating communication between primitive and definitive hepatoblasts across species (**Fig. 5b and 5c**).

To assess the causative role of PTN/MDK signaling in boundary organoids, we pharmacologically inhibited the pathway using AS1411, an NCL-specific aptamer known to suppress NCL phosphorylation and downstream PTN/MDK signaling^15^. Treatment with AS1411 resulted in a significant reduction in the area of *PROX1::mScarlet*⁺ cells, indicating a substantial impairment in hepatoblast expansion (**Fig. 5d**). These findings support a critical role for PTN/MDK signaling in promoting the proliferation within the boundary organoid model.

As highly proliferative embryonic tumor hepatoblastomas (HB) is characterized by the expression of primitive hepatoblast genes including *MDK*^16^, we further queried whether there is overlapping feature between primitive hepatoblast-derived tissues and HB. To test this, we’ve transplanted day 15 boundary organoid models and harvested the further developed graft at 2 months. Histological assessments confirmed presence of hepatic lineages: however, demonstrates mixed tissue formation comprised of the epithelial and mesenchymal tissue components including cartilaginous tissues (**Extended Data Fig. 7**). We noted these features are also present in mixed epithelial/mesenchymal type hepatoblastoma primary tissues (**Extended Data Fig. 7**). Primitive hepatoblasts exhibit unique molecular, spatial, and temporal profiles (**Fig 5e and 5f**) with a developmental potential for epithelial and mesenchymal hybrid tissues.

## Discussion

The transcription factors Sox2 and Cdx2 define the regional identity of the developing foregut and midgut, respectively^8^. Sox2 has been commonly used as a marker of the foregut, while Cdx2 is typically associated with the midgut with two boundaries forming around E8.75^12^. However, the expressions and roles of these transcription factors within the AIP lip around E8.0-E8.5, a region that gives rise to the liver, remain poorly understood. Interestingly, the expression patterns of Sox2 and Cdx2 do not align perfectly with the established boundaries of the foregut and midgut, complicating the interpretation of their roles in liver development^12^. This raises the possibility that Cdx2-positive endodermal cells could contribute to the hepatoblast population and studies in mouse embryo *ex vivo* cultures have suggested that midgut-derived cells are able to differentiate into hepatocytes, further implicating the potential of Cdx2-positive endodermal cells to differentiate into the hepatic lineage^17^. We observed scattered CDX2 expression in the lateral region of the AIP, which overlaps with the liver primordium^6^, provides spatial evidence supporting this developmental connection. Functionally, CDX2 has been shown to reprogram fibroblasts into hepatocytes by activating hepatic transcription factors such as *FoxA* and *Hnf4a*^18^, and its binding sites in the embryonic gut are enriched near genes involved in liver development^19^. Furthermore, the ability of hepatocytes to transdifferentiate into CDX2-positive intestinal epithelial cells through a hybrid epithelial/mesenchymal state resembling primitive hepatoblasts^20^ and that the hepatocyte differentiation are achieved from CDX2+ endodermal progenitors *in vitro*^21^. These, together with our findings, suggests that CDX2 progenitors may not be exclusive to intestinal fate but rather a developmentally plastic to liver fate.

Primitive hepatoblasts co-express hepatoblast markers (e.g., *TBX3, FGB, KRT8/18*), and mesenchymal markers (e.g., *VIM, PDGFRA, HLX*), while lacking mesothelial markers such as *PODOXL* and *MSLN*^22^. This hybrid transcriptional profile suggests that primitive hepatoblasts represent an intermediate state between hepatic epithelial progenitors and mesenchymal-like cells. Primitive hepatoblast-like subpopulations exhibiting mesenchymal characteristics have been identified in several prior studies. These include LIV2+ hepatoblasts that express Id3 and Mdk^1^, the hepatomesenchyme described at E10.5 in mouse liver^3^, and Id3+ hepatoblasts detected in both mouse (E11.5) and human (PCW5) fetal livers^4^. The reported frequency of these mesenchymal-like hepatoblasts ranges from 6–8%^4^ to as high as 57.6% (376 of 653 cells)^1^, with a consistent decline in abundance at later developmental stages^4^. Immunofluorescence analyses in these studies confirm co-expression of ID3^4^, GATA4, and VIM^3^ with conventional hepatoblast markers although the precise anatomical localization of these cells within the liver was not described. These findings support the existence of a transient mesenchymal-like hepatoblast population.

Primitive hepatoblasts express key mitogenic factors such as PTN and MDK, both of which have been implicated in promoting hepatocyte proliferation and regeneration. These factors have been reported to originate from a variety of sources within the liver including mesenchymal progenitor cells^23^, PODXL+ hepatic mesothelial cells^23^, and hepatic stellate cells^24^. Expression of MDK has also been identified as a maker of early hepatoblasts that downregulate over developmental time^25,26^, paralleling the decreasing proportion of primitive hepatoblasts. Both PTN and MDK have also been detected in hepatocytes, ductal plates, and biliary epithelial cells, further supporting their widespread involvement in hepatic growth^27,28^. The importance of these mitogenic cues is further underscored by studies showing that loss of the mesenchyme-associated transcription factor HLX results in liver hypoplasia^29^. The liver undergoes a phase of exponential growth at around E11.5 in mice. Mesenchymal cells are also thought to play a role, but their exact contribution has remained unclear. The identification of primitive hepatoblasts expressing *PTN* and *MDK* indicate that these hepatoblasts are a paracrine source of mitogens, supporting hepatoblast proliferation during this critical developmental window.

A primitive hepatoblast-like population has also been observed in a subset of hepatoblastomas, characterized by the co-expression of genes such as *ID3*^16,30^, *VIM*^31^, and *MDK*^16^ Hepatoblastoma, an embryonal liver tumor derived from hepatocellular precursors, recapitulates key features of early liver development and exhibits histological heterogeneity with both epithelial and mesenchymal elements, reflecting a reactivation or persistence of a primitive epithelial–mesenchymal hybrid state^32^. An "Intermediate" subpopulation with high *ID3* and *MDK* expression is enriched in patients with poor outcomes, supporting a model in which primitive hepatoblasts or their derivatives function as a cell of origin^16^. In co-culture, hepatoblastoma cells induced anti-inflammatory gene expression in macrophages, an effect reversed by MDK inhibition, implicating MDK in immune modulation^16^. Future studies incorporating immune cells are needed to further define its role in development and tumor progression.

Our findings support a model in which primitive hepatoblasts comprise a transient, spatially restricted, and molecularly distinct subset of liver progenitors. Characterized by a unique lineage origin, mesenchymal gene expression, and secretion of mitogenic factors, this population likely plays a specialized role in expanding the hepatoblast pool and coordinating early liver organogenesis. While their mesenchymal-like plasticity includes a capacity for cartilaginous differentiation, an observation relevant to recent human pluripotent stem cell-derived hepatocyte therapies^33^, the transient paracrine mitogenic signals, if controlled, may also offer an advantage in achieving the critical cell mass needed for effective regenerative interventions. Altogether, this population adds a new dimension to the developmental heterogeneity of the liver and underscores the importance of transient progenitor states in informing the design of safer and more robust liver regenerative therapies.

## Methods

### Lineage tracing

To trace the lineage of the early CDX2-positive cells, we employed tamoxifen inducible Cre-loxP and Dre-rox systems in mice. The transgene for the Cdx2^CreER^ through the Jackson laboratory (Jackson #022390^34^), was designed with a 9.5 kb promoter fragment from the 5’ flanking sequences identified through colon cancer cells^35^ that resulted in Cre-mediated recombination in distal ileal, cecal, colonic, and rectal epithelium but not in proximal intestine such as duodenum and jejunum^34^. Therefore, to facilitate lineage tracing of CDX2-positive domains in the gut tube of the proximal intestine, we developed a knock-in Cdx2^DreER^ mouse with P2A-DreERT2 cassette before the stop codon of the Cdx2 gene (**Extended Data Fig. 2a**). A comparison between the transgenic Cdx2^CreER^ and the knock-in Cdx2^DreER^ line was conducted using a dual-lineage tracing approach, combining Dre-rox and Cre-loxp strategies through the breeding of Rosa26-loxp-STOP-loxp-ZsGreen (Rosa26^LSLZsG^) and H11-rox-STOP-rox-tdTomato (H11^RSRtdT^) mice (**Extended Data Fig. 2a**). Notably, lineage tracing with transgenic Cdx2^CreER^ failed to induce recombination in the ventral midgut. However, recombination was observed in the dorsal midgut at E9.5, including the caudal half of the dorsal pancreas. In contrast, the Cdx2^DreER^ enabled lineage tracing of the anterior midgut irrespective of the dorsal-ventral orientation, although the recombination efficiency was lower in the dorsal midgut compared to Cdx2^CreER^.

#### Mouse embryos

All experiments involving mice were conducted in accordance with protocols approved by the Institutional Animal Care and Use Committee of Cincinnati Children’s Hospital Medical Center (CCHMC) (protocol IACUC2013-1028). Genotyping was performed by Transnetyx using real-time PCR.

Timed pregnancies were established by mating transgenic mice carrying inducible CreERT2 or DreERT2 alleles with appropriate reporter strains. The day of vaginal plug detection was designated as embryonic day 0.5 (E0.5). To induce recombination, either 4-hydroxytamoxifen (4-OHT; Sigma-Aldrich) or tamoxifien (Sigma-Aldrich) was used. 4OHT was dissolved in Kolliphor EL (Sigma) containing 25% ethanol and administered via intraperitoneal injection at a dose of 66 ug/g body weight^36^, and tamoxifen was dissolved in corn oil at 20mg/mL and administered at a dose of 75ug/g body weight. Injections were performed between E7.5 and E10.5, depending on the experimental design.

Embryos were collected at the time of interest ranging from E8.5 to E18.5. Pregnant females were euthanized in accordance with institutional animal care and use guidelines, and uteri were dissected in cold PBS. Embryos were isolated under a stereomicroscope. Fluorescence was assessed when applicable, and embryos were fixed in 4% paraformaldehyde (PFA) at 4 °C overnight. Following fixation, embryos were washed in PBS and processed for downstream applications such as whole-mount immunostaining, cryosectioning, or imaging.

### Maintenance of hPSCs

Human PSC lines were maintained under feeder-free conditions as previously described. Briefly, undifferentiated PSCs were cultured in mTeSR1 medium (STEMCELL Technologies) or StemFit medium (Ajinomoto Co., Japan) on plates coated with either Matrigel Growth Factor Reduced (Corning; 1:30 dilution) or Matrix-511 (Nippi, Japan; 0.25 µg/cm²). Cultures were maintained at 37 °C in a humidified incubator with 5% CO and 95% air.

### Differentiation of hPSCs into AG and PG spheroids

Differentiation of human PSCs into definitive endoderm was performed as described previously with minor modifications^37^. Briefly, iPSCs were dissociated using Accutase (Thermo Fisher Scientific) and seeded at 150,000 cells/mL on Matrigel-coated plates. The differentiation medium was changed daily as follows:

Day 1: RPMI 1640 with 100 ng/mL activin A (R&D Systems) and 50 ng/mL BMP4 (R&D Systems)

Day 2: RPMI 1640 with 100 ng/mL activin A and 0.2% defined fetal bovine serum (dFBS)

Day 3: RPMI 1640 with 100 ng/mL activin A and 2% dFBS From days 4 to 7, cells were cultured in gut growth medium [Advanced DMEM/F12 supplemented with 15 mM HEPES, 2 mM L-glutamine, penicillin–streptomycin, B27 (Life Technologies), and N2 (Gibco)]:

Anterior gut induction: with 200 ng/mL noggin (R&D Systems), 500 ng/mL FGF4 (R&D Systems), and 2 µM CHIR99021 (Stemgent)

Posterior gut induction: with 500 ng/mL FGF4 and 3 µM CHIR99021

On the day of spheroid formation (typically day 7), anterior or posterior gut cells were dissociated into single cells using a 1:9 mixture of TrypLE Express 10X (Life Technologies) and 0.05% trypsin at 37 °C. The cells were then centrifuged at 300G for 3 minutes and resuspended in gut growth medium supplemented with 10 µM Y-27632 (Tocris Bioscience). Cell suspensions were plated in 96-well ultra-low attachment V-bottom plates (S-BIO) at a density of 10,000 cells per well and incubated for 24 hours to allow spheroid formation. The next day, individual PG spheroids were transferred to the well containing AG spheroid gut and co–cultured for 24 additional hours to generate fused anterior–posterior (AP) boundary spheroids. These AP boundary organoids were either embedded to Matrigel the following day or kept in floating culture in ultra-low 96 well V-bottom plates (S-BIO) in gut growth medium.

### Kidney capsule transplantation

Day 15 AP boundary organoids generated from D7 AG and D7 PG spherodis were transplanted into the subcapsule of the kidney in immune-deficient NOD.Cg-*Prkdc^scid^ ll2rg^tm1Wji^* Tg(CMV-IL3,CSF2,KITLG)1Eav/MloySzJ (NSGS) mice at 8 weeks of age and harvested after 2 months. All experiments were performed under the approval of the Institutional Animal Care and Use Committee of Cincinnati Children’s Hospital Medical Center (protocol number IACUC2023-1028).

### Whole-mount immunostaining of organoids, spheroids, and embryos

Organoids embedded in Matrigel were released by gentle pipetting and washed with PBS. After centrifugation at 200 × g for 3 minutes (repeated if necessary), samples were transferred to 1.5 mL tubes and fixed in 4% paraformaldehyde (PFA) at 4 °C overnight. Spheroids and embryos were collected directly into 1.5 mL tubes, allowed to settle, and washed with PBS prior to overnight fixation in 4% PFA at 4 °C. Following fixation, all samples were washed five times in PBS and permeabilized in 0.1% Tween-20 for 30 minutes. After washing with PBST (0.1% Triton X-100 in PBS), samples were blocked in 1% BSA and 0.3% Triton X-100 in PBS for 1 hour at room temperature.

Samples were incubated with primary antibodies diluted in blocking buffer (200 µL per tube or 150 µL in spot plates for visible samples) at 4 °C overnight (or up to 2–3 days for larger samples) on a rocker. Antibody solutions were centrifuged at 13,000 × g for 10 minutes prior to use to reduce nonspecific staining. The following day, samples were washed five times in PBST for at least 1 hour each. Fluorophore-conjugated donkey secondary antibodies (1:500 dilution in blocking buffer) and DAPI were added and incubated overnight at 4 °C in the dark. After secondary incubation, samples were washed five additional times in PBST and mounted in Fluoroshield with DAPI. For tissue clearing, samples were embedded in 4% low-melting agarose at 37 °C, then immersed in refractive index matching solution (RIMS; iohexol-based, pH 7.5) for ≥12 hours before imaging. H&E staining and immunohistochemistry

### Multi-plex immunofluorescent staining

Frozen tissue sections were thawed at room temperature for 10-20 minutes, while paraffin sections were de-paraffinized and subjected to antigen retrieval. Sections were then rinsed in PBS for 10 minutes (repeated 2-3 times) and permeabilized with 0.1% Tween-20 in PBS for 30 minutes. Blocking was performed using 1% BSA and 0.3% Triton X-100 in PBS for 1-2 hours. Primary antibodies, diluted in blocking buffer, were applied and incubated overnight at 4°C. The following day, slides were washed and secondary antibodies (1:500 dilution) were applied, followed by overnight incubation at 4°C in the dark. After additional washing, slides were mounted with Fluoroshield and coverslipped.

For multiplex imaging, we utilized Alexa Fluor 405 (or DAPI), Alexa Fluor 488, Alexa Fluor 555, Alexa Fluor 594, Alexa Fluor 647 (Invitrogen), Brilliant Violet 480 (Jackson ImmunoResearch), and CF750 (Biotium). Fluorophores were carefully selected to minimize spectral overlap, particularly avoiding combinations such as Alexa Fluor 555 and 594 with tdTomato/mScarlet. Due to the large size of BV proteins, they were primarily used on tissue sections rather than wholemounts to reduce background fluorescence. Imaging was conducted using a Nikon AXR Inverted Confocal Microscope equipped with 8 high-power solid-state diode lasers (405, 445, 488, 514, 567, 594, 647, and 730 nm), and appropriate filters were selected to minimize spectral overlap.

### RNA isolation and RT–qPCR

Total RNA was extracted using the RNeasy Mini Kit (Qiagen). Reverse transcription was carried out using the High-Capacity cDNA Reverse Transcription Kit (Applied Biosystems) following the manufacturer’s instructions. Quantitative PCR (qPCR) was performed using TaqMan Gene Expression Primers (Thermo Fisher Scientific) on a QuantStudio 5 Real-Time PCR System (Thermo Fisher Scientific).

### RNA sequencing

Whole-transcriptome RNA sequencing was performed on hPSC-derived AG, PG, and boundary organoids dissected from the boundary region. Library preparation and sequencing were carried out by Novogene (China) using the Illumina NovaSeq S4 platform with 150 bp paired-end reads, generating ∼20 million reads per sample. Raw reads were quality-filtered to remove adapter sequences, reads with >10% unknown bases (N), and low-quality reads (>50% of bases with quality scores ≤5). Clean FASTQ files were aligned to the human hg38 reference genome using the Computational Suite for Bioinformaticians and Biologists (CSBB v3.0) to obtain raw transcript counts.

Log2 counts per million (CPM), normalized by the Trimmed Mean of M-values (TMM) method, were used for differential expression analysis with the Gene Expression Analysis Kit^38^ and Gene Set Enrichment Analysis^39^. Genes with a false discovery rate (FDR) < 0.05, fold change >1.5, and expression >0.5 CPM in at least one sample were considered significantly differentially expressed.

### Single-Cell Transcriptomic Data for the developing liver in mouse and human

Publicly available scRNA-seq datasets were used to investigate early liver development in mouse and human. Mouse embryonic liver data were obtained from ArrayExpress (E-MTAB-9334), and human fetal liver data from Wesley et al. (E-MTAB-8210, E-MTAB-7189, E-MTAB-7407). Sequencing reads were processed using the Cell Ranger Software Suite and aligned to the GRCh38 reference genome to generate UMI count matrices.

Data preprocessing and quality control were performed using Seurat v5 in R (v4.4.0). Data were normalized using Seurat’s default global-scaling method. Dimensionality reduction was performed using the top 30 principal components, followed by UMAP for visualization. Clustering and differential expression analysis were also conducted using Seurat.

Cell–cell communication was analyzed using CellChat. Annotated cell types were used to infer ligand–receptor interactions, compute communication probabilities, and visualize aggregated interaction networks using default parameters.

### Hepatocyte and Hepatoblast Isolation

Primary hepatocytes were isolated from anesthetized adult mice using a two-step liver perfusion protocol, as previously described^40^. Liver Perfusion Medium (LPM; Thermo Fisher Scientific) and Liver Digest Medium (LDM; Thermo Fisher Scientific) were prewarmed to 45 °C for 30 minutes. Percoll-HBSS (1×) and wash medium were prepared and maintained at 4 °C. Following abdominal disinfection, a 24 G catheter was inserted into the inferior vena cava (IVC), and 45 mL of LPM was perfused through the liver over 5 minutes, followed by a second perfusion with 45 mL of LDM. The liver was excised, minced in ice-cold wash medium, and filtered through a 70 μm cell strainer into Percoll-HBSS. The resulting cell suspension was centrifuged at 300 × g for 10 minutes at 4 °C, and the pellet was washed and resuspended for further analysis.

Embryonic hepatoblasts were isolated by dissecting livers from embryos, followed by rinsing with ice-cold 2% FBS in PBS (2%FBS/PBS). The tissue was then incubated and mechanically dissociated with micro scissors in LPM at 37 °C for 10 minutes. After centrifugation at 300 × g for 3 minutes, the pellet was digested in LDM at 37 °C for 15– 30 minutes. Dissociated cells were filtered through a 70 μm cell strainer (Corning, NY), centrifuged at 300 × g for 5 minutes, and treated with RBC lysis buffer (eBioscience) for 15 minutes on ice to remove red blood cells. Cells were then washed and resuspended in 2%FBS/PBS for downstream analysis.

### Flow Cytometry

For flow cytometric analysis, cultured cells were dissociated using a 1:9 mixture of TrypLE Express 10× (Life Technologies) and 0.05% trypsin at 37 °C for 15–30 minutes. Dissociated hepatoblasts and hepatocytes were resuspended in 2%FBS/PBS and incubated with anti-Fc receptor antibody for 20 minutes on ice to block nonspecific binding. Cells were stained with primary antibodies for 30 minutes, washed in 2%FBS/PBS, and centrifuged at 300 × g (50 × g for adult hepatocytes) for 5 minutes. Secondary antibody staining was performed for 20 minutes on ice. To exclude dead cells, samples were incubated with Zombie UV viability dye (BioLegend) in PBS without FBS at room temperature prior to analysis. For intracellular staining, cells were fixed, permeabilized, and stained using the BD Pharmingen™ Transcription Factor Buffer Set (BD Biosciences) according to the manufacturer’s protocol.

### CUT & RUN

CUT & RUN was performed as described previously^41^. Briefly, concanavalin A-coated beads (20 µL per reaction; Bangs Laboratories, BP531) were washed twice with Binding Buffer (20 mM HEPES-KOH [pH 7.9], 10 mM KCl, 1 mM CaCl□, 1 mM MnCl□). Live cells were harvested, counted, and resuspended (100,000 for H3K4me3; 250,000 for H3K27me3) in Wash Buffer (20 mM HEPES-NaOH [pH 7.5], 150 mM NaCl, 0.5 mM spermidine, 1× EDTA-free protease inhibitor cocktail). Beads were added to the cells and incubated with rotation at room temperature for 10–15 minutes. The cell-bead complexes were then resuspended in 100 µL of Antibody Buffer (2 mM EDTA in Digitonin Buffer, composed of 0.02% digitonin in Wash Buffer), and incubated overnight at 4 °C with rotation in the presence of primary antibodies. Following two washes with Digitonin Buffer, samples were incubated with 700 ng/mL of pA-MNase for 1 hour at 4 °C with rotation. Samples were then chilled on a CoolBlock for at least 10 minutes. Digestion was initiated by adding 2 µL of 0.1 M CaCl□, followed by incubation at 0 °C for 30 minutes. Reactions were stopped by adding 100 µL of 2× Stop Buffer (340 mM NaCl, 20 mM EDTA, 4 mM EGTA, 0.02% digitonin, 0.05 mg/mL RNase A, 0.05 mg/mL glycogen). CUT&RUN fragments were released by incubating at 37 °C for 30 minutes with shaking (300 rpm). Beads were pelleted by centrifugation (16,000×g, 5 minutes, 4 °C) and removed using a magnetic stand. The supernatant was treated with 0.1% SDS and 0.25 mg/mL Proteinase K for 1 hour at 37 °C. DNA was then extracted via phenol-chloroform and ethanol precipitation. DNA concentration was quantified using a Quantus Fluorometer (Promega)

### Human hepatoblastoma (HB) samples

We have collected background liver and tumor samples from patients with HB under conditions appropriate for characterization with institutional review board approval (IRB # 2016-9497) and informed patient consent. Primary tumor and background liver were used to examine histological features, as described previously^42^.

## Acknowledgments

The authors thank A. Kodaka for scientific illustration materials; Y.C. Hu and C. Mayhew for generating the PROX1::mScarlet-edited hPSC line; M. Kofron for guidance with immunostaining, confocal imaging, and image analysis; and S. Rankin for critically reading the manuscript. We are grateful to all members of the Takebe, Zorn, Helmrath, and Wells laboratories for their support and feedback. S. Rankin for critically reading the manuscript. We also acknowledge the CCHMC Bio-Imaging and Analysis Facility (RRID:SCR_022628), Integrated Pathology Research Facility (RRID:SCR_022637), Pluripotent Stem Cell Facility (RRID:SCR_022634), Transgenic Animal and Genome Editing Facility (RRID:SCR_022642), Single Cell Genomics Facility (RRID:SCR_022653), Genomics Sequencing Facility (RRID:SCR_022630), Research Flow Cytometry Facility (RRID:SCR_022635) and Veterinary Services Facility.

This work was supported by the Cincinnati Children’s Center for Pediatric Genomics Trainee Award and a Japan Society for the Promotion of Science (JSPS) Overseas Research Fellowship to K.I., who is also a New York Stem Cell Foundation (NYSCF)– Druckenmiller Fellow. Additional support was provided by NIH grants P30 DK078392 and R01GM143161 to M.I.; This work was also supported by Cincinnati Children’s Research Foundation CURE grant, the Falk Transformational Award Program, NYSCF, NIH Director’s New Innovator Award (DP2 DK128799-01) and R01DK135478 to T.T. This work was also supported by an NIH grant UG3/UH3 DK119982, Cincinnati Center for Autoimmune Liver Disease Fellowship Award, PHS Grant P30 DK078392 (Integrative Morphology Core and Pluripotent Stem Cell and Organoid Core) of the Digestive Disease Research Core Center in Cincinnati, Takeda Science Foundation Award, Mitsubishi Foundation Award, Cannon Foundation Award and by Japan Agency for Medical Research and Development (AMED) under grant numbers: JP24bm1223006, JP24ym0126809, JP21bm0404045, JP18fk0210037, JP18bm0704025, JP23gm1610005, JP23fk0210106, JP24gm1210012, JP24fk0210150 and JP21fk0210060, JST Moonshot: JPMJMS2022 and JPMJMS2033, JSPS KAKENHI: JP18H02800, 19K22416, 21H04822 and World Premier International Research Center Initiative (WPI) PRIMe, MEXT, Japan.

## Author Contributions

K.I., H.K, and T.T. conceived the study, K.I and T.T. designed the experiments, K.I. wrote original draft of the manuscript, M.M, M.I, J.M.W, A.M.Z, and T.T. reviewed and edited the manuscript, K.I., H.K, Y.M. and M.K. performed *in vitro* experiments, K.I., K.K., and C.S. performed animal maintenance, embryo collection, section, staining and analysis, K.I., K.G. and A.B. analyzed hepatoblastoma histology, K.I., M.G. H.W.L., K.T. and M.I. performed and analyzed CUT&RUN analysis, K.I., H.A.R., N.S. and K.T. performed and analyzed RNA-seq and scRNA-seq analysis.

## Figure Legends

**Extended Data Figure 1. Spatial distribution of CDX2 during hepatic specification.**

**a.** Wholemount immunostaining of E8.5 embryos showing SOX2 (green), CDX2 (red), and PROX1 (blue) along the anterior-posterior axis. CDX2 is detected in scattered cells near the anterior intestinal portal lip overlapping with PROX1+ cells.

**b–c**. Wholemount immunostaining of E8.75 (b), and E9.5 and E10.5 (c) embryos showing the hepato-biliary-pancreatic region. PROX1+ cells (green) emerge at the boundary between SOX2+ (blue) and CDX2+ (magenta) domains and quantification of SOX2+ and CDX2+ cell overlap with PROX1+ region in the ventral and dorsal PROX1+ buds.

**Extended Data Figure 2. Generation and validation of Cdx2^DreER^ lineage tracing model.**

**a.** Schematic of the knock-in strategy introducing a P2A-DreERT2 cassette immediately before the stop codon of Cdx2 and wholemount staining of CDX2^CreER^ CDX2^DreER^ dual lineage tracing at E9.5.

**b.** Confocal imaging confirmed tdTomato labeling in tamoxifen-induced embryos expressing SOX2^CreER^, CDX2^DreER^, or FOXA2^CreER^. Overlapping tdTomato expression was observed in CDX2^DreER^ and FOXA2^CreER^ lineage tracing models.

**c.** Immunostaining demonstrated that CDX2^DreER^ -labeled cells co-expressed hepatoblast markers including KRT8/18, FGB, DLK1, and TBX3, mesenchymal marker VIM, and epithelial marker E-cadherin at the ventral edge of the developing liver. Additionally, some tdTomato cells overlapped with the endothelial marker CD31.

**Extended Data Figure 3. Persistence of Cdx2-derived hepatoblasts into late gestation and adulthood.**

**a.** Flowcytometry analysis through E10.5 to E14.5 detects the increase of tdTomato+ cells within the hematopoietic (CD31-CD45+) lineage and endothelial (CD31+) lineage within the liver.

**b.** At E18.5, tdTomato+ hepatocytes (E-cadherin+ Epcam-) were evenly distributed in all lobes. **c**.Isolation of adult hepatocytes revealed ∼1% of ASGR+ cells retained tdTomato signal, confirming a small persistent Cdx2-derived hepatocyte population.

**Extended Data Figure 4. Characterization of PROX1::mScarlet reporter line and hepatic induction in human boundary organoids.**

**a.** Schematic of the knock-in strategy introducing an mScarlet cassette into human embryonic stem (ES) cells.

**b.** Karyotype analysis confirmed genomic stability following the knock-in procedure.

**Extended Data Figure 5. Clustering analysis of scRNA-seq of day 12 human boundary organoids**

**a.** Clustering analysis of scRNA-seq data from day 12 boundary organoids identified primitive hepatoblast-like clusters, as well as a distinct definitive endoderm-like cluster.

**Extended Data Figure 6. Epigenetic regulation of RA pathways in boundary organoids.**

**a.** Gene ontology analysis revealed significantly stronger repression of retinoic acid (RA) signaling in anterior gut (AG), marked by enrichment of the repressive histone modification H3K27me3 in RA response-related biological process pathways.

**Extended Data Figure 7. Histological comparison of transplanted organoids and hepatoblastoma**.

**a.** Grafts of day 15 boundary organoids collected after 3 months show mixed tissue formation, including hepatic, mesenchymal, and cartilage-like structures by H&E staining. Similar features are seen in primary hepatoblastoma tissue from three independent patients.

