## Supplementary Figures for "Primitive Hepatoblasts Driving Early Liver Development"

Extended Data Figure 1. Spatial distribution of CDX2 during hepatic specification.

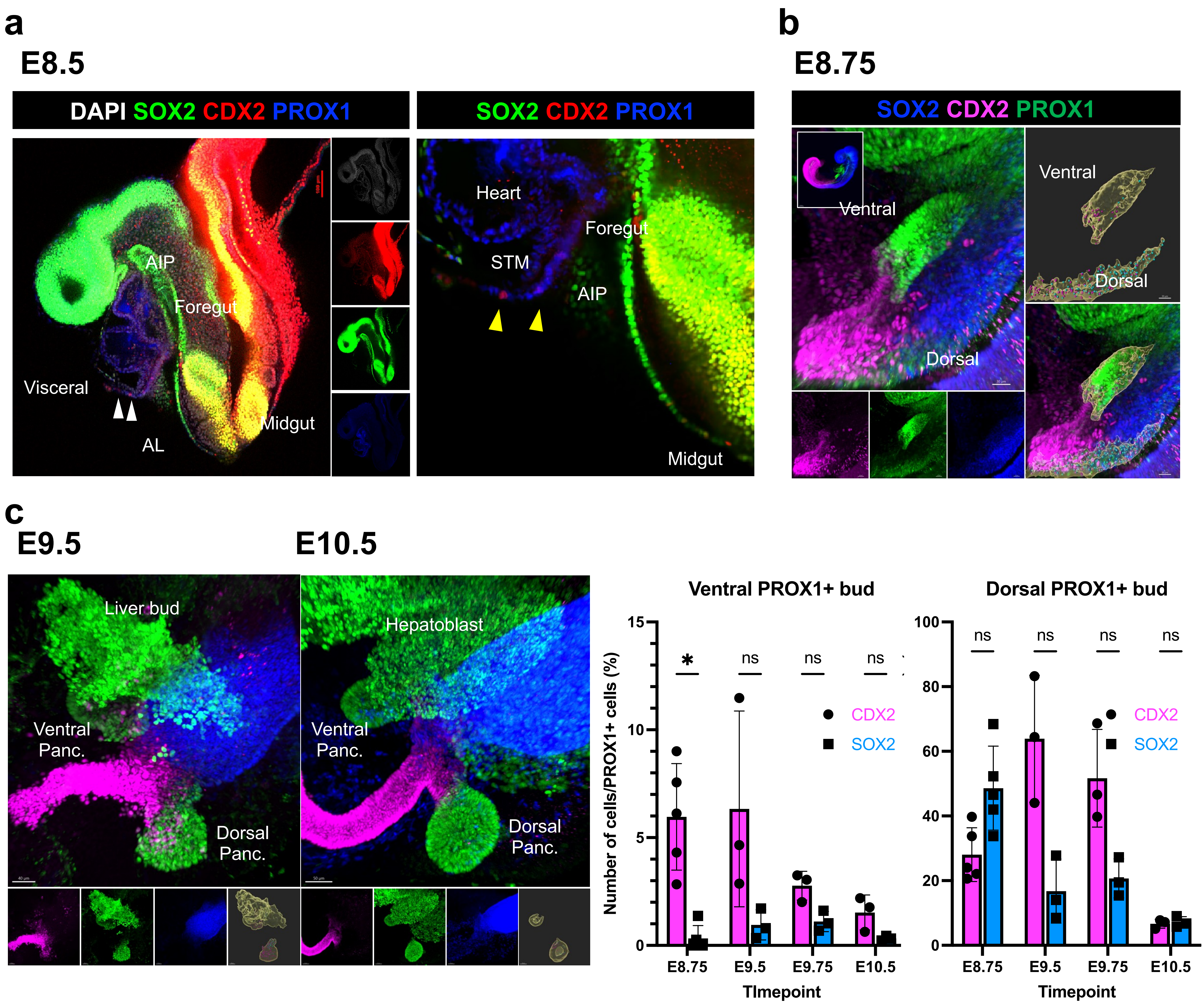

Extended Data Figure 2. Generation and validation of Cdx2DreER lineage tracing model.

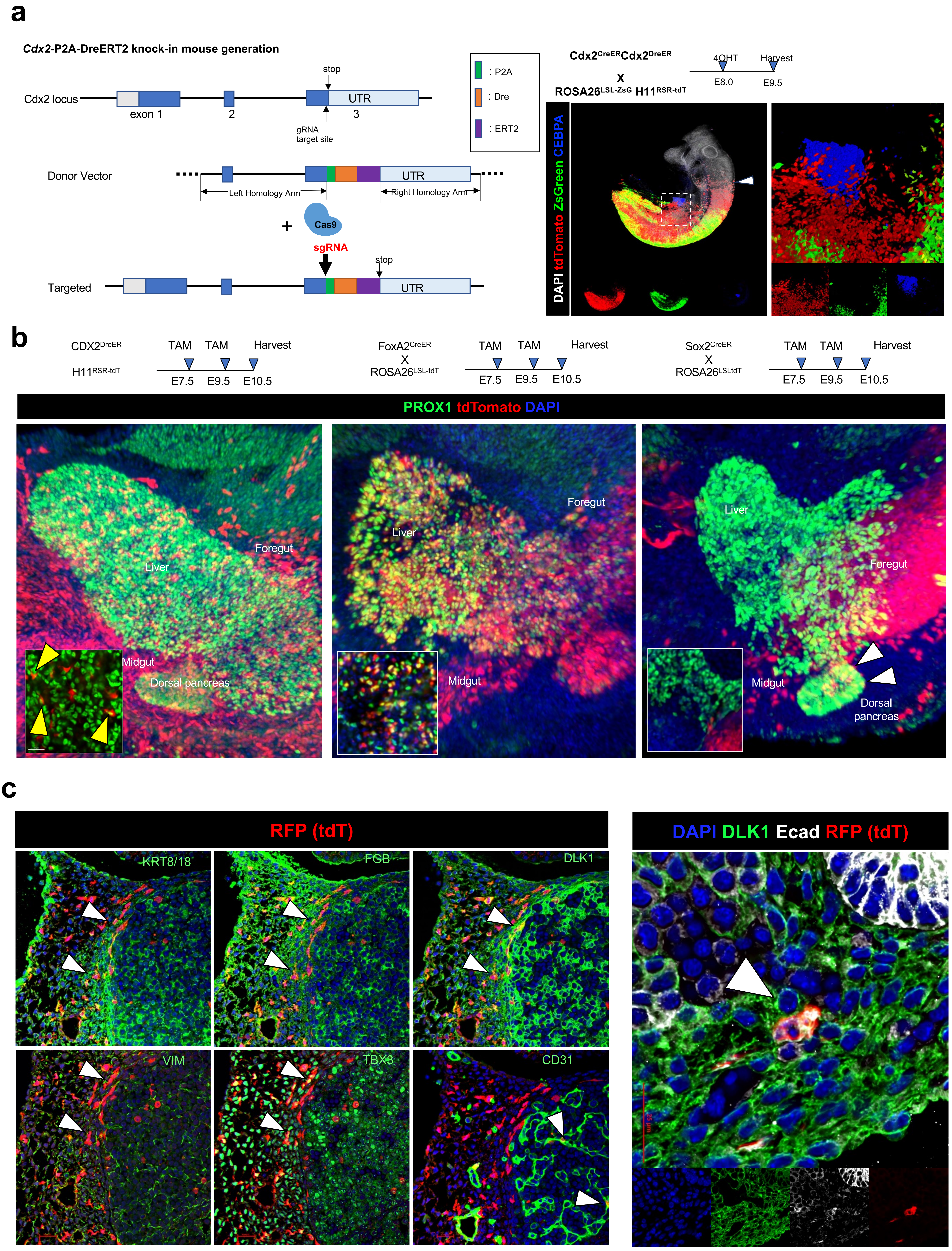

Extended Data Figure 3. Persistence of Cdx2-derived hepatoblasts into late gestation and adulthood.

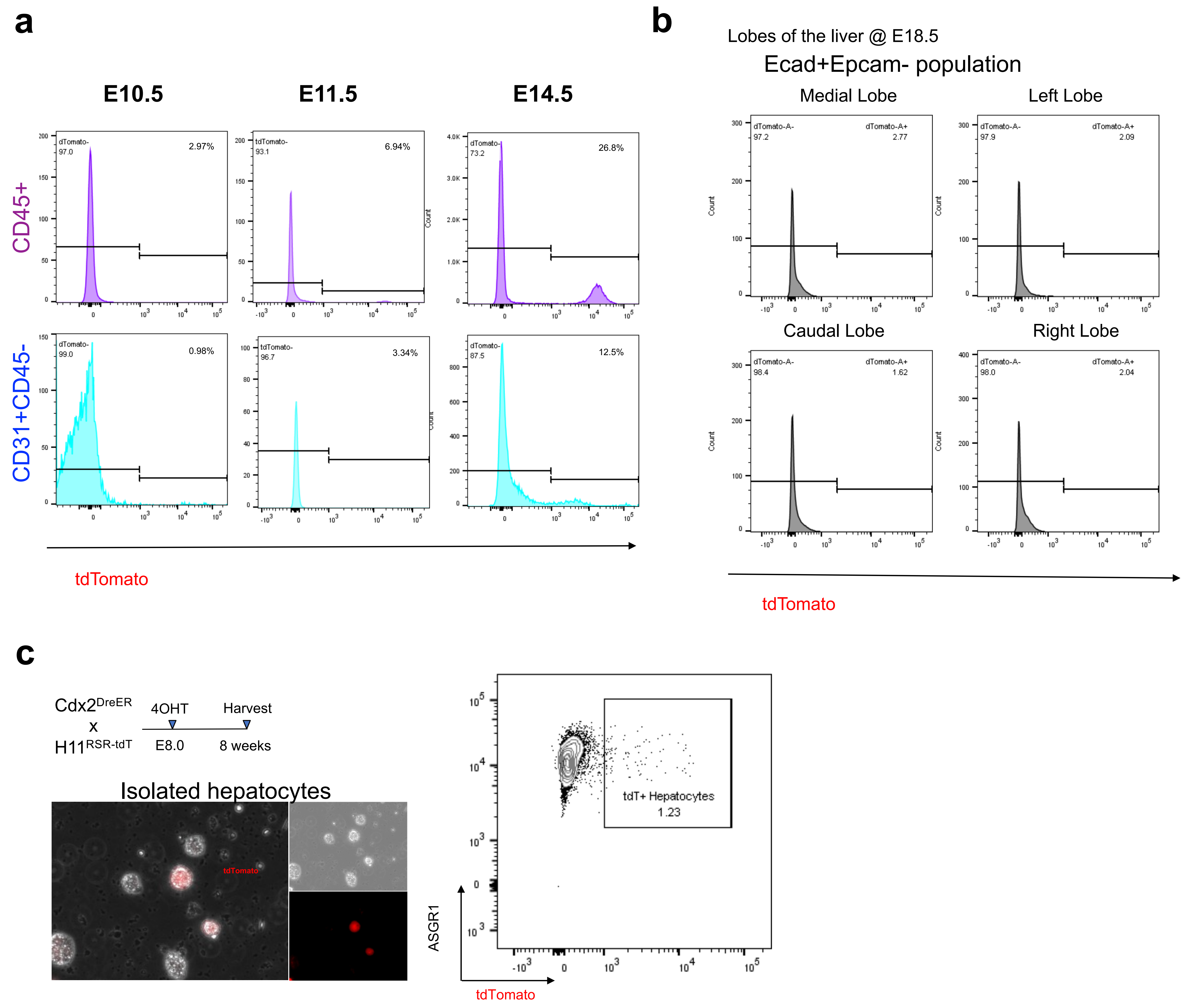

Extended Data Figure 4. Characterization of PROX1::mScarlet reporter line and hepatic induction in human boundary organoids.

a

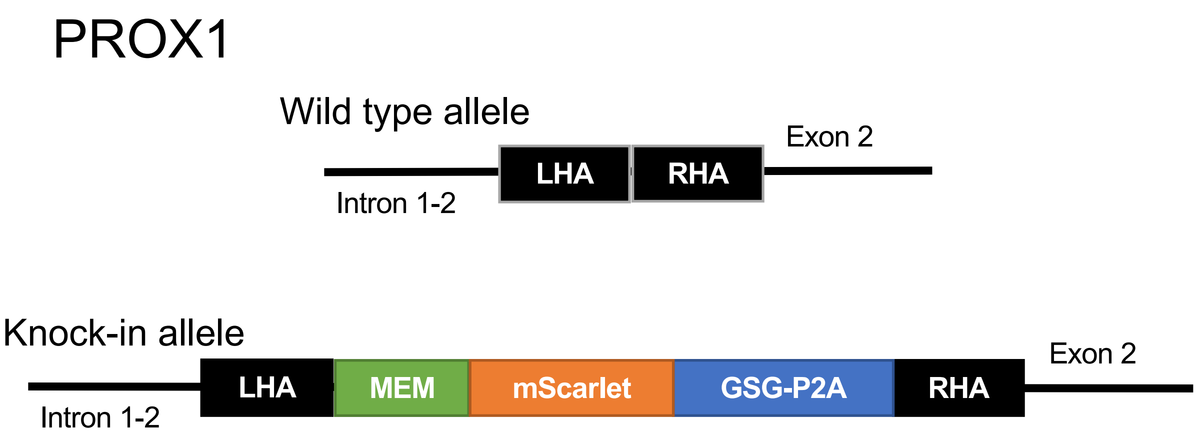


b

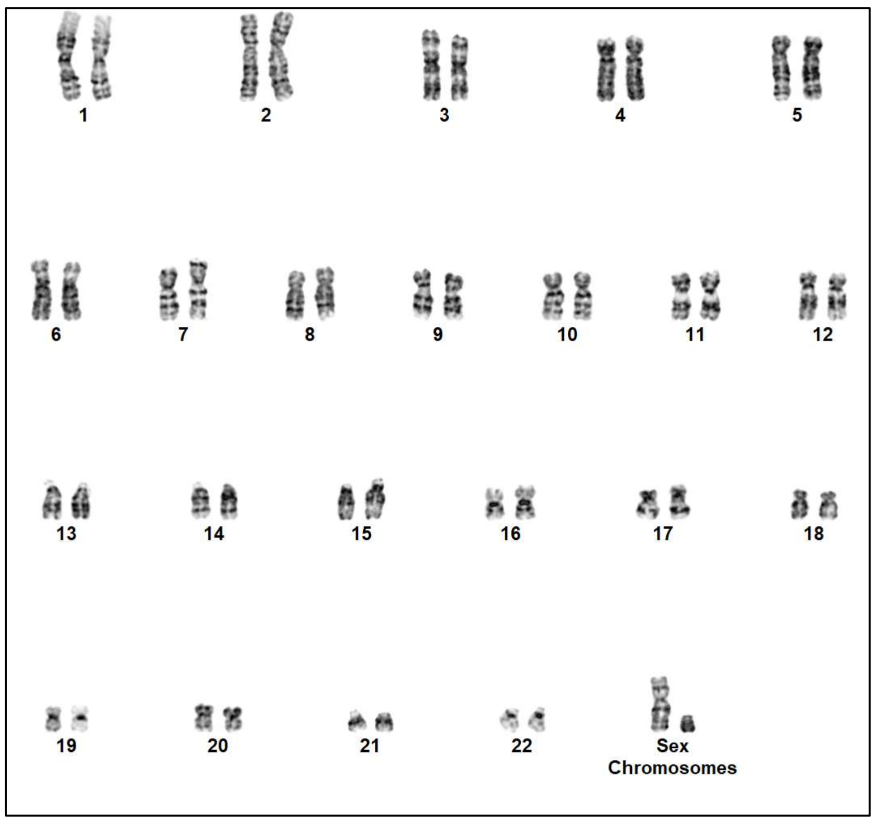

Extended Data Figure 5. Clustering analysis of scRNAseq of day 12 human boundary organoids.

**a**

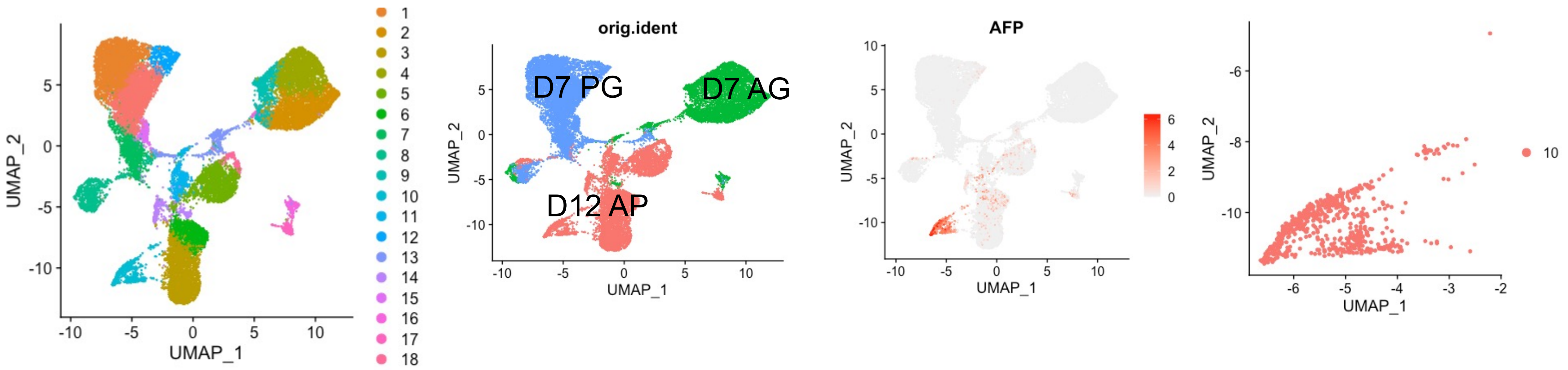

Extended Data Figure 6. Epigenetic regulation of RA pathways in boundary organoids.

**a**

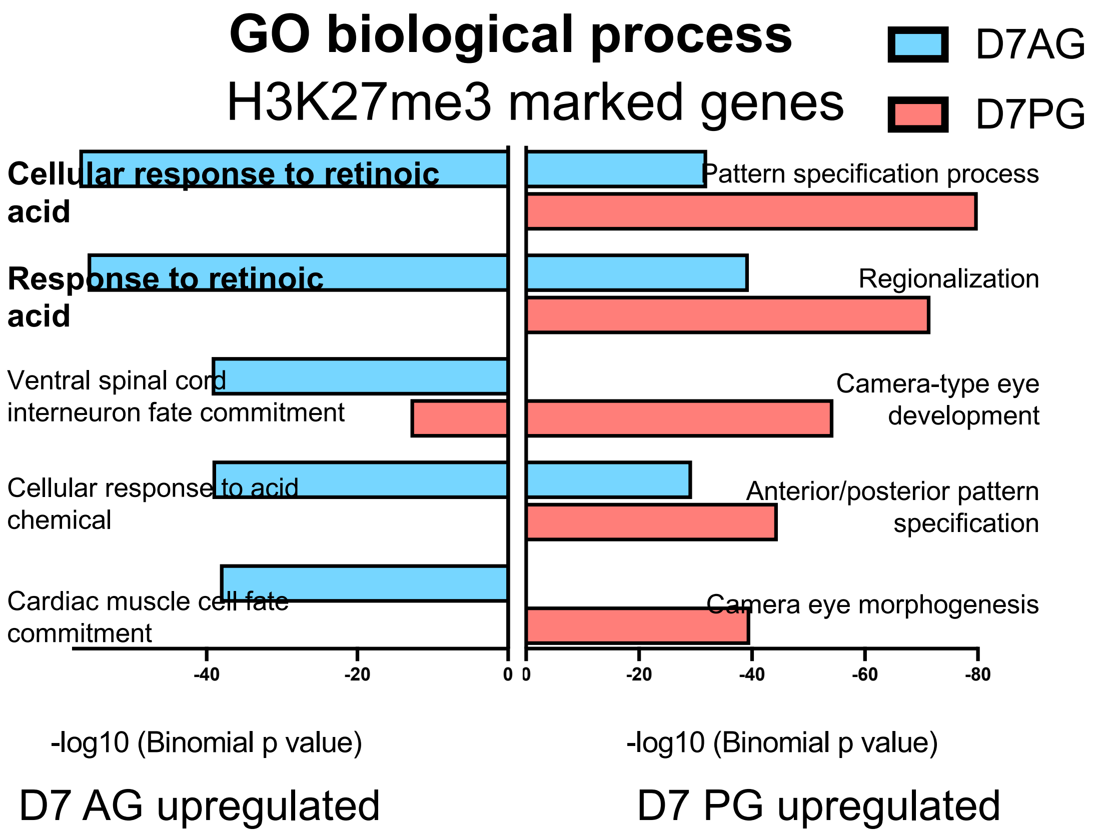

Extended Data Figure 7. Histological comparison of transplanted organoids and hepatoblastoma

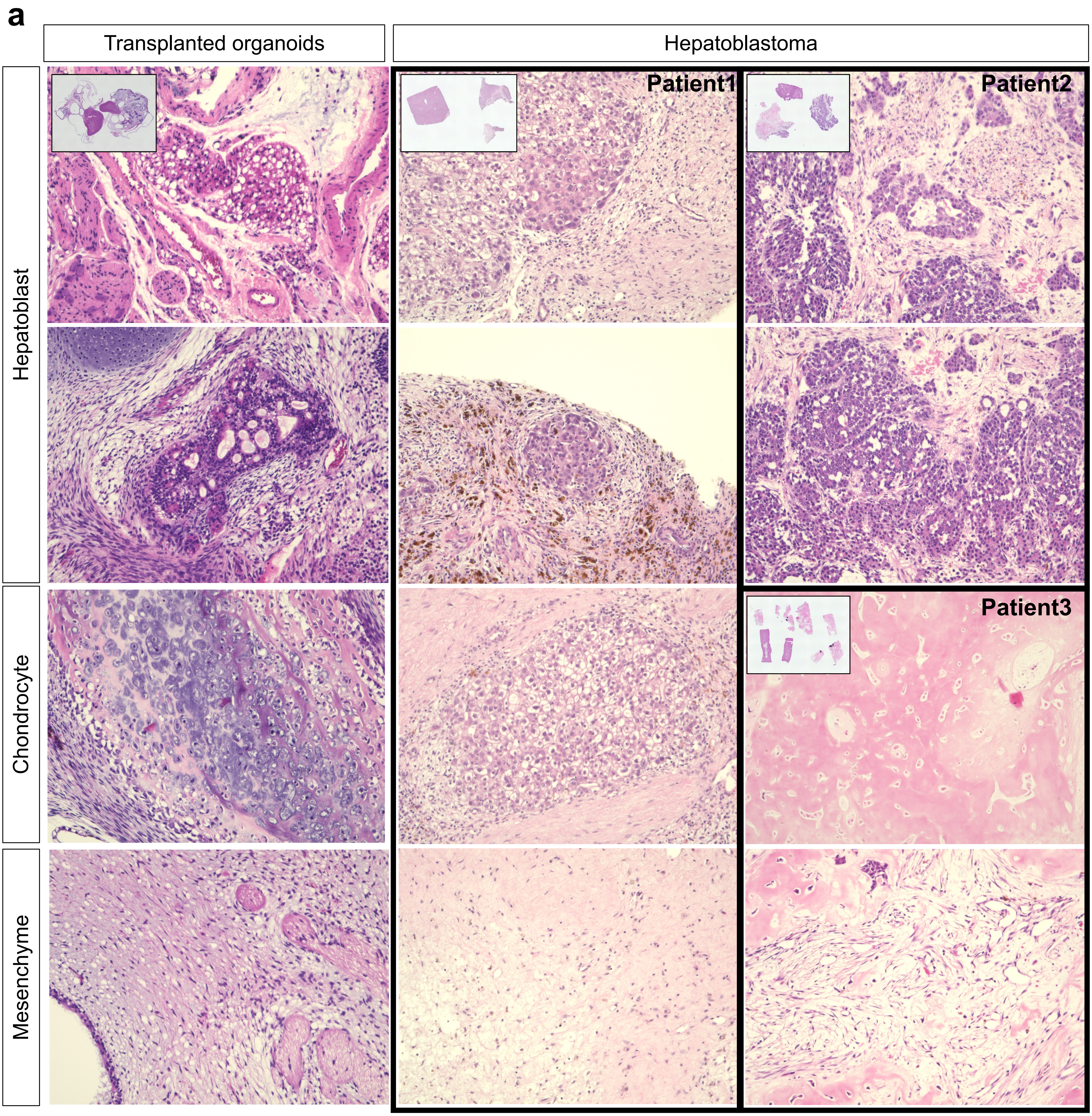
